# A Stochastic Neural Mass Model for Cortical Beta Bursts in Parkinson’s Disease

**DOI:** 10.64898/2026.08.10.743870

**Authors:** James Ross, Brian Skelly, Zelekha Seedat, Matthew Brookes, Stephen Coombes, Áine Byrne

**Affiliations:** School of Mathematical Sciences, University of Nottingham, Nottingham, UK; School of Mathematics and Statistics, University College Dublin, Dublin, Ireland; Sir Peter Mansfield Imaging Centre, University of Nottingham, Nottingham, UK

## Abstract

Beta-band (13–30 Hz) oscillations are increasingly understood to occur as transient “bursts” rather than sustained rhythms, with altered burst dynamics, specifically increased duration and power alongside reduced burst rates, in patients with Parkinson’s disease (PD). In this study, we utilise resting state magnetoencephalography (MEG) data from healthy adults to quantify the temporal fluctuations in the beta-band, and examine the distributions of burst statistics. We then fit a stochastic next-generation neural mass model to these empirical statistics using a Genetic Algorithm. Systematic parameter sweeps reveal that reducing background drive to excitatory and inhibitory neuronal populations reproduces the altered burst statistics observed in PD. Crucially, we show that strengthening synaptic coupling can counteract these deficits and restore healthy bursting dynamics. Together, this work establishes a computational framework linking cellular-level mechanisms to macroscale burst statistics, and highlights potential targets for therapeutic neuromodulation in movement disorders.

**Author summary:** Brain activity is comprised of rhythmic electrical patterns called “brain waves.” Traditionally, these waves were viewed as smooth and continuous, but recent evidence reveals that they actually occur in brief, intense bursts. In conditions such as Parkinson’s disease, these bursts become altered—lasting longer, growing stronger, and occurring less frequently. In this study, we developed a mathematical model of brain tissue to understand what drives these burst patterns. Using real brain scans from healthy human volunteers, we tuned our model with an optimisation algorithm until its simulated bursts closely matched real human brain activity. We then systematically varied the model’s settings to investigate how abnormal bursting arises in disease. We discovered that reducing the background signals to the brain cells reproduces the burst alterations seen in Parkinson’s disease. Importantly, our simulations showed that strengthening the connections between brain cells can counteract this deficit, restoring healthy burst patterns. By connecting microscopic cell properties to whole-brain rhythms, our work offers new insights into how movement disorders disrupt brain networks and highlights potential cellular targets to guide future brain stimulation therapies or medications.

## Introduction

Neural oscillations have long fascinated experimental and theoretical neuroscientists alike. Believed to arise from the synchronised activity of vast ensembles of neurons, these rhythmic oscillations are readily observed using non-invasive techniques such as electroencephalography (EEG) or magnetoencephalography (MEG). They occur at a range of different frequencies and are categorised into distinct frequency bands, each associated with different brain states and functions. Beta-band oscillations (13-30 Hz) are of particular interest due to abnormal activity in the beta band being linked to a variety of brain diseases and disorders, such as Parkinson’s disease and schizophrenia.

Classically, beta oscillations were thought to be sustained oscillations that varied in amplitude over time due to certain tasks being performed. However, this picture was formed by looking at trial averaged data and more recent studies investigating non-averaged data suggests that beta oscillations occur as transient increases in power in the beta frequency range, rather than smooth oscillations [1–3]. These increases in power have been coined *beta bursts*. Several recent studies have investigated the functional role of beta bursts and shown that modulation of trial-averaged beta power reflect event-related increases and decreases of the burst probabilities [4–6]. In particular, Little *et al.* showed that periods of event-related synchronisation observed in motor cortex post movement were dominated by high amplitude beta bursts, with a strong correlation between the timing of the bursts and movement initiation [7]. Burst statistics have also been shown to be modulated by cognitive load. Rodriguez-Larios and Haegens showed that, during a working memory task, the burst power and duration decreased with memory load and during memory manipulation, while the burst rate increased [8].

Pathological beta activity in the cortico-basal ganglia loop is commonly associated with Parkinson’s disease (PD) [9] [10]. A number of recent studies examining beta bursts in the cortex observed an increase in burst power and duration, coupled with a decrease in the number of bursts per second. When comparing PD patients on and off medication to healthy controls, Vinding *et al* [11] reported a reduction in the mean burst rate by 5-17% in patients with PD. They note that this is the most affected of the three beta burst characteristics in PD patients. In a similar study, Pauls *et al* [12] also reported a reduction in the burst rate for PD patients when their DBS was off, and found that the beta burst duration and the beta burst power are increased.

There appears to be conflicting evidence on the impact of PD on beta activity as the characteristics of the beta burst vary substantially throughout the cortico-basal ganglia loop. For example, in the subthalamic nucleus (STN) the burst rate is reported to increase in patients off medication versus on medication [13]. However, an increase in burst duration and power appears to be consistent across brain areas and measurement techniques [13–15]. In this study, we employ a mathematical model of cortical cells, and as such, focus on experimental findings from cortical data.

From a mathematical modelling perspective, stochastic neural mass models are able to support bursting like behaviour. Powanwe and Longtin employed a stochastic two population Wilson-Cowan model to investigate bursting behaviour in the gamma frequency band (*>* 30 Hz) and developed a statistical averaging method to link model parameters to burst duration and envelope [16]. However, the Wilson–Cowan model is phenomenological in nature and fails to account for some important aspects of neurodynamics, such as neuronal synchronisation. Here, we consider the next generation neural mass model, which has many of the features of the Wilson–Cowan model, albeit with the added advantage of a dynamic description for the evolution of synchrony and a direct link to an underlying spiking network [17]. Given the success of the next generation neural mass model in replicating movement induced changes in the beta rhythm [18], it is a natural candidate for studying beta bursts. In our 2020 review article, we showed that adding stochasticity to the next-generation neural mass model resulted in bursting behaviour similar to that seen in MEG data [19]. We presented an example *single trial* spectrogram that showed clear bursts in beta-band power, as well as *trial-averaged* spectrogram across many simulations (Figure 5.1 of [19]), albeit with no attempt to to match the burst characteristics to real data.

Burst detection methods have typically employed some form of threshold detection where the time series data (from EEG or MEG recordings) is filtered by frequency band and an amplitude threshold is used to determine when the bursting occurs. More recently, hidden Markov models (HMM) have been employed to identify bursting states in MEG time series data. This technique provides a more objective way of detecting burst states by looking at specific spectral patterns rather than just amplitude in a single frequency band [20, 21]. In this work, we simulate a stochastic next-generation neural mass model and employ the HMM model to identify bursting states in simulation data, as well as in healthy control MEG data. We compare the burst statistics of the MEG data and the simulation data, and employ a Genetic Algorithm (GA) to optimise the model parameters such that the model burst statistics closely match those from the real data. We then perform a number of parameter manipulations to better understand the importance of each parameter in the generation of bursting behaviour, with the hope of shedding a light on the generation of pathological bursting behaviour in diseases such as Parkinson’s disease and proposing mechanisms to restore healthy bursting behaviour.

## Results

### Modelling fitting

In our previous work [19], we reported that a stochastic next-generation neural mass model could produce beta bursts, but did not investigate the burst characteristics (duration, power, rate). Here, we employ a hidden markov model (HMM) to detect beta bursts in both the MEG data and the output of the stochastic neural mass model. We then calculated the average burst duration and burst power, as well as the average number of bursts per second (burst rate) for each trial. Figure 1B shows the distribution of each statistic across trials. The real data are shown in blue, and the simulated data for the unoptimised parameters from [19] are shown in orange. The mean burst duration for the model data is around 50 ms, while the mean of the real data is around 250 ms. The distribution is also much tighter for the model data compared to the real data. For the burst power, the model data and real data have similar means, but the width of the distribution is much greater for the real data. The model averages around 4 bursts per second, while in the real data we see only around 1 burst per second. The model does, however, have a reasonably good fit on the spread of the distribution.

**Fig 1.**
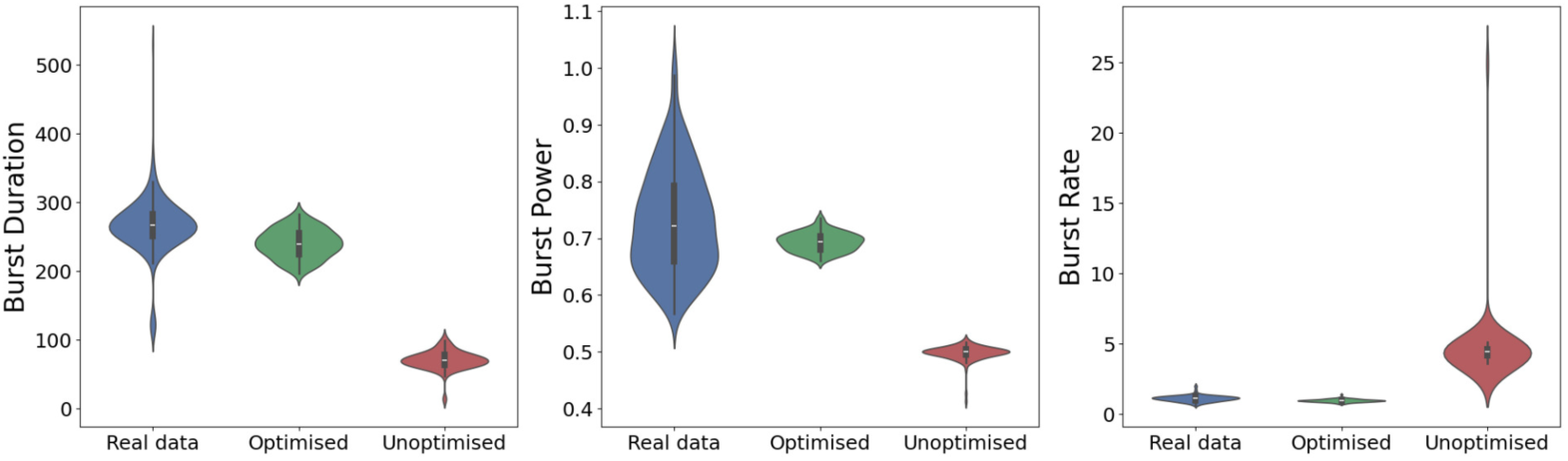
Burst statistics. Distributions of the burst statistics; duration (left), power (middle), rate (right), from the MEG data (blue) and the neural mass model with optimised parameters with a fitness of 1.9 (green) and non optimised parameters with a fitness of 20 (red).

We then employ a Genetic Algorithm (GA) to fit the model parameters to replicate the real bursting activity (see *Materials and Methods* for details of the model and fitting procedure). Comparing the burst statistics of the model with optimised parameters to the real data, we find a much improved fit (Fig. 1 green, fitness score ∼ 1.9). The mean burst duration is now significantly closer to the mean burst duration in the real data, and the spread of the distribution is also closer to the real data. The mean power is again similar to the value for the real data, but unlike the unoptimised model data, the spread of the data has increased and the distribution is much more similar to that of the real data. For the number of bursts we see an excellent fit between the model data and the real data, the GA has helped the model to get much closer to the mean value of around 1 burst per second. In all statistics here, the GA has improved the fit to the real data, with the most significant improvement being in the burst rate.

### Manipulating burst statistics

Next we explore how changing the model parameters affects the burst statistics. In general, we find that when the burst duration increases, the power also increases, while the burst rate decreases.

Increasing the mean background drive of the excitatory population from the optimised value *η_E_* = 2 results in a reduction in the burst power, but the burst duration and burst rate remain more or less unchanged (Fig. 2). While reducing *η_E_*leads to an increase in the burst duration and burst power, with a decrease in the burst rate. There are no obvious trends in the spread of the distributions, with the long narrow tails on some distributions being the result of single outliers. Looking at the mean background drive of the inhibitory population *η_I_* , we see that the burst power peaks close to the optimised value of *η_I_* = 1.04. Interestingly, the spread of the distribution increases as *η_I_*increases but not as it decreases. The burst duration and burst rate also have local maxima/minima around this point, although the increases and decreases are less pronounced. The width of both distributions increases as *η_I_* increases or decreases away from the optimised value.

**Fig 2.**
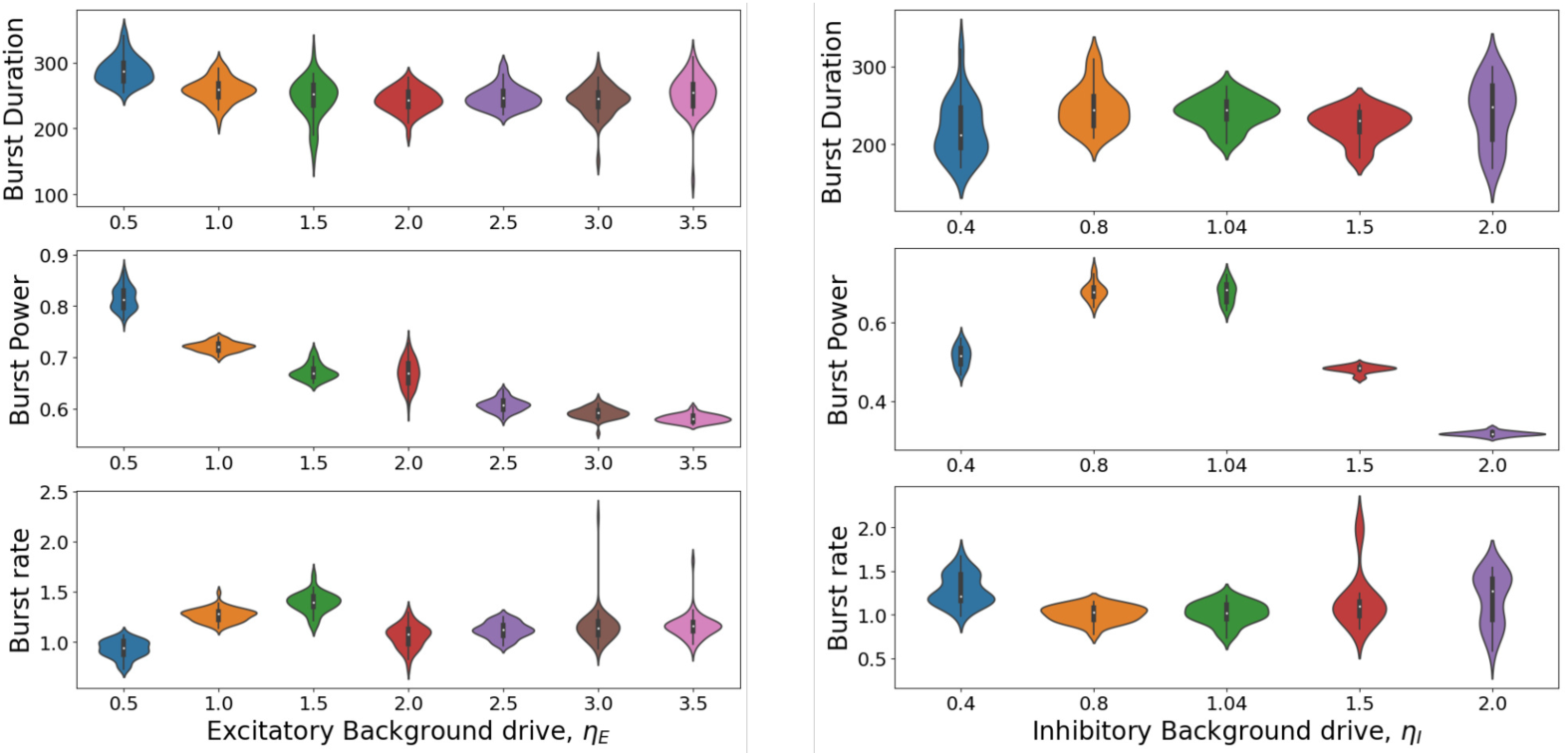
Effects of changing the mean background drive. Distributions of the burst statistics: duration (top), power (middle) and rate (bottom), as the mean background drive to the excitatory population *η_E_* (left) and inhibitory population *η_I_*(right). Each violin shows the distribution of the mean burst duration/power/rate across 25 trial. All other parameters are set to the optimised values.

The next parameter of interest is the level of heterogeneity (Fig. 3). Increasing the level of heterogeneity of the excitatory population from the optimised value Δ*_E_* = 0.2 leads to a moderate increase in the burst rate, coupled with minor increases and decreases in the burst duration and power, respectively. Reducing Δ*_E_* from 0.2 results in an increase in the duration and power of the bursts, coupled with a decrease in the burst rate. The width of the burst power distribution decreases when Δ*_E_* is changed from the optimised value of 0.2, decreasing substantially for higher values of Δ*_E_*. There is very little change in the distribution widths for both the burst duration and burst rate. Now, looking at the heterogeneity of the inhibitory population, modifying Δ*_I_* from the optimized value of 0.2 leads to a reduction in the burst power, with a greater reduction for increases in Δ*_I_* . The width of the distribution also reduces for increases and decreases in Δ*_I_* , with decreases in Δ*_I_* resulting in very narrow distributions. The behaviour of the burst duration and burst rate is less straightforward. The burst duration initially decreases and then increases as Δ*_I_* is increased, while the burst rate initially increases and then stabilises with a mean value of around 1.3. Decreasing Δ*_I_* results in shorter and more frequent bursts, with a wide distribution of both characteristics for Δ*_I_* = 0.15, which appears bimodal. The width of the burst duration distribution increases for large Δ*_I_* ,

**Fig 3.**
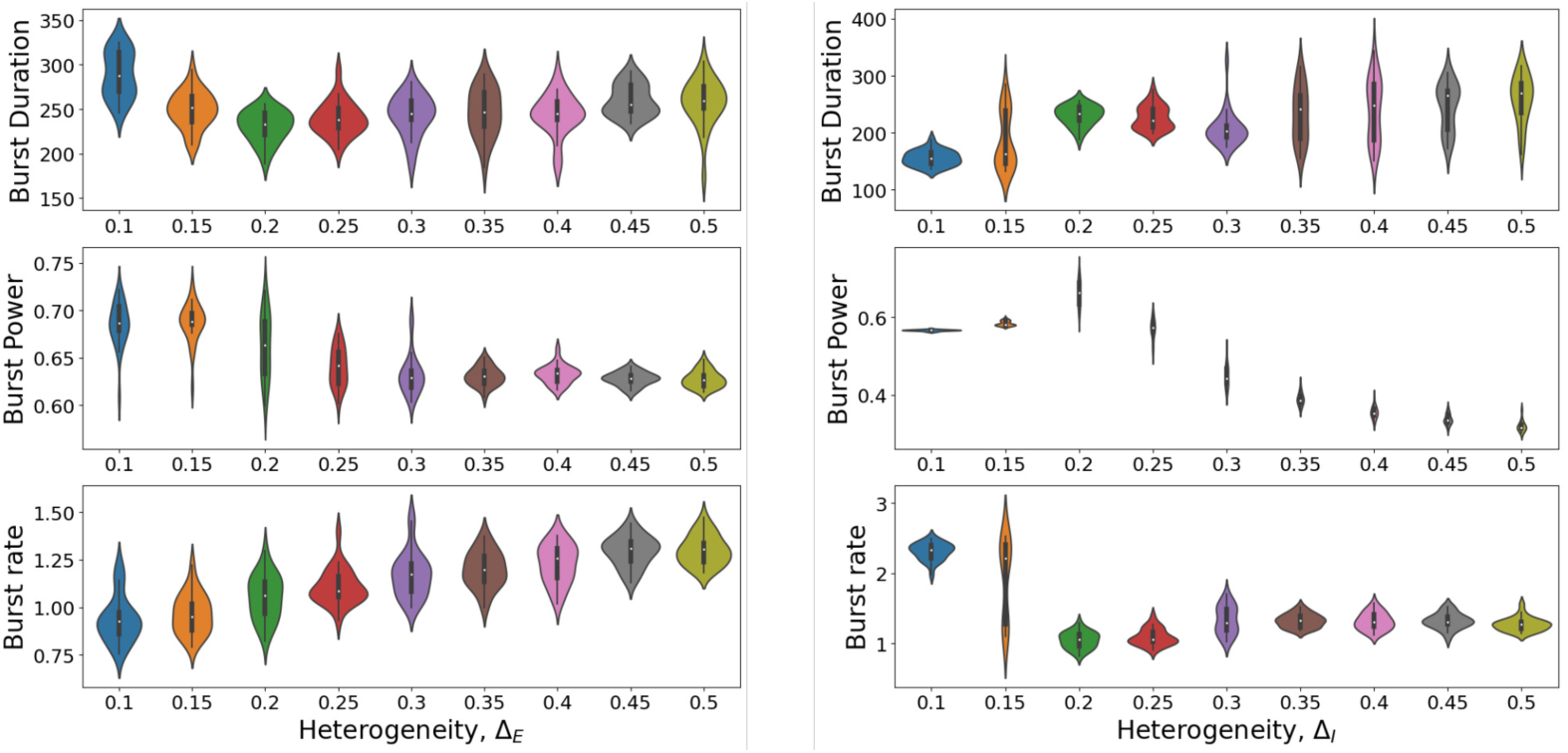
Effects of changing the level of heterogeneity. Distributions of the burst statistics, duration (top), power (middle) and rate (bottom), as the level of heterogeneity in the excitatory population *η_E_*(left) and inhibitory population *η_I_*(right). As in Fig. 2, each violin shows the distribution of the means across 25 trial. All other parameters are set to the optimised values.

We found that changing the gap junction coupling strength between excitatory neurons 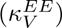 did not greatly affect the burst statistics. The strength of the connections between excitatory and inhibitory neurons (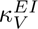 and 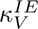) did change the burst statistics, but not by a huge amount. The gap junction coupling strength between inhibitory neurons (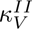 was found to be close to optimal in terms of maximising burst duration and power, while minimising burst rate (see Fig. S1 in the Supplemental Material). The final parameters we studied were the synaptic time constants which were found to have little effect on the burst statistics (see Fig. S2 in the Supplemental Material).

Noting that reducing background drive to the excitatory population *η_E_* resulted in Parkinsonian-like burst characteristics, less frequent, longer, higher power bursts, we examine this range more closely while also varying background drive to the inhibitory population *η_I_*(Fig. 4). When the background drive to the inhibitory population *η_I_*is low, we see short, low-power bursts occurring more regularly. As *η_I_* is increased, the bursts become longer, the power increases, and the burst rate decreases. We observe a maximum in the burst power and a minimum in the burst rate in the range *η_I_ ∼* 0.6 − 0.9 for all values of *η_E_*, with the extent of the increase/decrease increasing ever so slightly with *η_E_*. The mean burst duration also increases in this region; however, we also observe increases in the region *η_E_ >* 1.1*, η*_1_ *>* 1.1, where the burst power decreases and the burst rate increases. Guided by Vinding et al.’s findings [11], we define Parkinsonian parameter values to be those that result in a 5-17% reduction in the mean burst rate, which also result in an increased burst duration and power. These parameter regions are highlighted in red.

**Fig 4.**
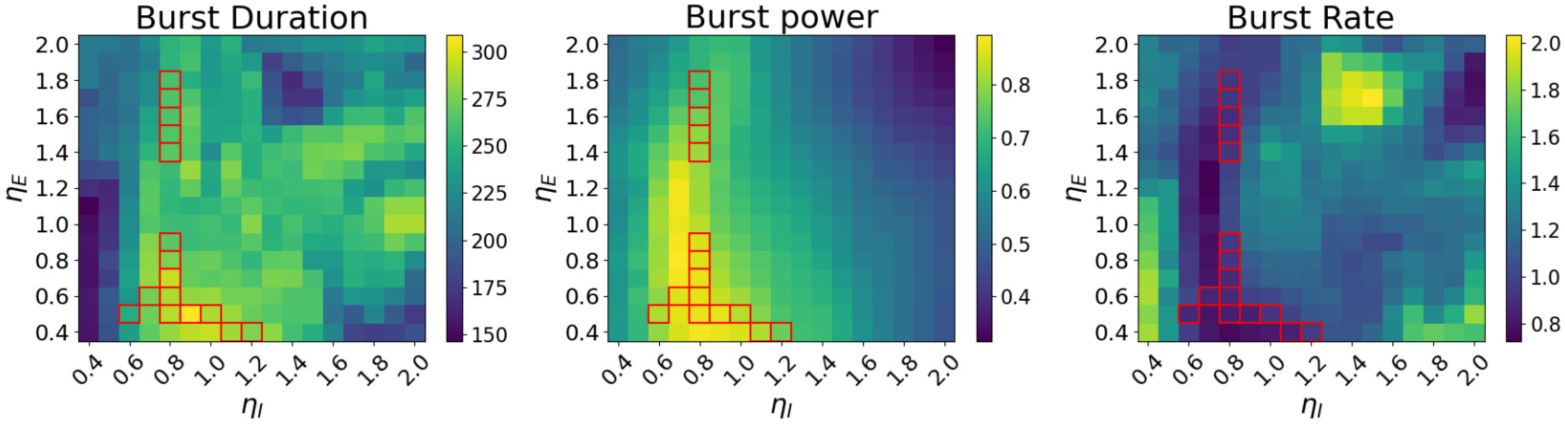
Pathological beta burst. Colour plots of the mean burst duration (left), power (middle), and rate (right), as the background drive to the excitatory *η_E_* and inhibitory populations *η_I_*are varied. Parameter sets that result in Parkinsonian burst activity (5-17% reduction in the mean burst rate) are highlighted with red squares. The results from the optimised parameter values of *η_E_*= 2.0 and *η_I_* = 1.04 (healthy controls) are found in the middle of the top row for each plot. All other parameters are set to the optimised values.

Finally, we explore whether alternative parameter manipulations can restore healthy beta activity. Many pharmacological interventions for neurological diseases are designed to impact synaptic transmission. Hence, we study the effect of manipulating the synaptic coupling strengths. As such interventions do not typically distinguish between postsynaptic neuron types, we scale both excitatory coupling strengths by the same factor 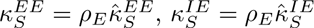 and both inhibitory coupling strengths by a different factor 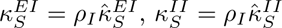, where 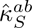 is the original optimised parameter value. We decrease the background drive to the excitatory population to *η_E_* = 0.5, which resulted in Parkinsonian burst activity with an 19.5% increase in burst duration, a 20.3% increase in burst power, and a 7.6% decrease in burst rate, and examine how scaling the synaptic coupling strengths affects the burst statistics (Fig. 5). Starting at *ρ_E_* = 1, *ρ_I_* = 1 (optimised coupling strengths), we see that increasing the excitatory coupling, *ρ_E_*, decreases the burst power and increases the burst rate, while the burst duration increases ever so slightly. Reducing *ρ_E_* has the opposite effect. Now examining the inhibitory coupling, we find that increasing *ρ_I_* increases the burst power, but the burst duration and rate do not change very much. Decreasing *ρ_I_*decreases both the burst power and the burst rate, while the burst duration is increased somewhat. To decrease the burst duration substantially, it is necessary to decrease both *ρ_E_* and *ρ_I_* , but this further decreases the burst rate into the Parkinsonian range. Given that burst rate is the most commonly studied statistic in Parkinson’s disease, we focus on the yellow region in the burst rate panel where the burst rate is restored to a healthy level (relative value *≈* 1). The burst power is also reduced, but the burst duration remains unchanged. Interestingly, we find that increasing *ρ_I_* as well as *ρ_E_* further increased the burst rate. Combining these observations, we conclude that strengthening both the excitatory synapses and inhibitory synapses is the most promising approach to restore healthy beta activity in the cortex, with a larger increase for the excitatory synapses compared to the inhibitory synapses.

**Fig 5.**
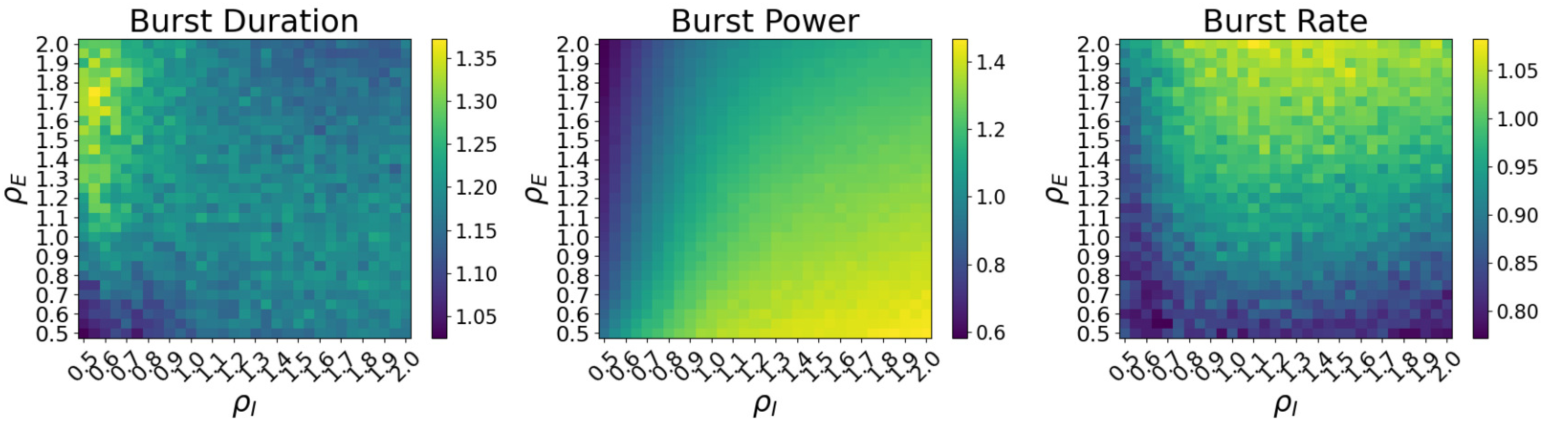
Restoration of healthy activity. Colour plots of the mean burst duration (left), power (middle), and rate (right), relative to their mean values in the healthy case, as the strength of the synaptic connections are varied. Restoration of the bursting behaviour to a health range occurs when the relative statistics are equal to 1. The mean background drives are set to the pathological values *η_E_*= 0.5 and *η_I_* = 1.04. All other parameters are set to the optimised values.

To demonstrate the robustness of our findings, we performed a similar analysis for an alternative set of Parkinsonian parameter values, namely *η_E_* = 0.8 and *η_I_* = 0.8, which resulted in an 14.6% increase in burst duration, a 26.9% increase in burst power, and a 8.1% decrease in burst rate (see Supplemental Material Fig. S3). Again, we found that increasing *ρ_E_*decreases the burst power and increases the burst rate, with minimal effect on the burst duration. Decreasing *ρ_I_* initially increases the burst duration, but decreasing further results in a decrease. This is similar for the burst rate, but we have the opposite dependence, the burst rate initially decreases, followed by an increase for larger decreases in *ρ_I_* . The burst power decreases monotonically with *ρ_I_* . To restore a healthy burst rate and power in this case, strengthening just the excitatory synapses may be sufficient, but strengthening the inhibitory synapses also will still lead to a reduction in the burst power and an increase in the burst rate. The two sets of colour plots show similar trends, with the area *ρ_E_∼* 0.5 − 1.2 and *ρ_I_ ∼* 0.7 − 1.5 in this plot closely matching the full area of the other.

## Discussion

In this work, we tune a stochastic next generation neural mass model to produce realistic beta bursting activity. As in previously published studies of both MEG and EEG data, we employ a hidden Markov model to detect and characterise the bursts. We optimise the model parameters using a Genetic Algorithm to match the burst statistics to MEG bursting data for healthy controls. Using the optimised parameters as a baseline, we perform a number of parameter sweeps to understand how abnormal bursting behaviour may arise in diseases such as Parkinson’s disease. We find that decreasing the background drive for both excitatory and inhibitory neurons (*η^E^*_0_ and *η^I^*_0_) leads to less frequent, longer, higher power bursts, as seen in PD patients. We also found that increasing the strength of the excitatory synapses restored the pathological bursting behaviour to a health range.

### Findings on Parkinson’s disease

The primary neurological degradation in PD is seen in the basal ganglia, and results in increased basal ganglia activity. This increased basal ganglia activity over inhibits the thalamus, which projects to the cortex and results in reduced excitability/external drive to cortical areas [22–24]. This aligns with our findings that reducing the background drive, which can be thought of as input from elsewhere in the brain, results in Parkinsonian bursting activity.

Common pharmacological treatments for neurological drugs target neurotransmitter release and synaptic response, which is why we choose to manipulate the synaptic coupling strengths in an attempt to restore healthy bursting activity from the Parkinsonian state, with decreased external drive to both populations. One problem with pharmacological interventions for neurological disorders is the blood-brain barrier. Not only is it difficult to design drugs that can effectively cross this barrier, but the drug is diffused across the entire brain and high doses are required to achieve therapeutic concentration levels [25–27]. For example, amantadine, a glutamate NMDA receptor channel blocker, is widely prescribed for reducing dyskinesia in Parkinson’s disease. In a 1995 study, Kornhuber *et al.* collected brain tissue from autopsies of patients treated with amantadin-sulfate during their lifetime and reported that amantadine concentrations were homogeneously distributed in brain areas [28]. Hence, glutamate was suppressed across the basal ganglia, cerebellum, etc, as well as the cortex. Designing targeted pharmacological treatments could prove challenging, but there is a growing body of research into strategies to transport drugs across the blood–brain barrier [29]. Another alternative for local manipulations of the synaptic activity is optogenetic stimulation or transcranial magnetic stimulation TMS.

There is a growing body of evidence that the cortex reorganises to mitigate the reduced thalamic input, with TMS showing improvements in motor symptoms [30–32]. This is also in line with our findings that manipulating the synaptic coupling strengths modifies the burst statistics. Fennelly *et al.* recently included synchrony-driven plasticity in a similar next generation neural mass model [33]. An interesting follow-up study would be to employ the plasticity framework of Fennelly *et al.* to study the effect of TMS on this network, and determine the optimal TMS stimulation protocols to restore healthy beta-band activity in the cortex.

### Limitations of the Genetic Algorithm

While the Genetic Algorithm (GA) significantly improved model fitness, it was ultimately limited in its ability to find highly optimised parameters. A primary factor for this is the simplification of fitting a single-node model to whole-brain data, which fails to capture the complex effects of inter-regional neural coupling. Future iterations could address this limitation by fitting a more sophisticated stochastic network model to the MEG data. Although running multiple Monte Carlo trials for a network-level model is computationally intensive, this approach is becoming increasingly feasible with the continued advancement and availability of GPU-accelerated computing.

Several algorithmic enhancements could also be implemented to improve the optimisation process. For example, replacing the hidden Markov model (HMM) currently used for burst detection with a computationally cheaper thresholding method [34] would accelerate the GA. Addressing memory constraints—either by optimizing the codebase or utilizing larger RAM capacities—would allow for increased trial and population sizes. Additionally, replacing fixed mutation strengths and crossover probabilities with adaptive alternatives could improve algorithmic flexibility; for instance, dynamically increasing mutation rates during periods of stagnation would help the GA escape local minima (See [35] for an array of comparisons between static and adaptive mutation). Finally, the fitness function itself could be refined. Rather than relying on twelve summary statistics, it may be more informative to directly compare the shapes of the target distributions using Earth Mover’s Distance (EMD) [36]. This would reduce the evaluation to just three metrics (EMD for burst power, duration, and rate), which could further mitigate the issue of local optima.

In summary, this study demonstrates that a stochastic next-generation neural mass model can successfully replicate the characteristic beta burst statistics observed in resting-state human MEG data. Parameter sweeps revealed that reducing the background drive generates the prolonged, higher-power, and less frequent beta bursts associated with Parkinson’s disease. Furthermore, our findings indicate that strengthening synaptic coupling, particularly excitatory connections, can counteract this deficit and restore healthy bursting dynamics. These findings provide important mechanistic insight into the network dynamics underlying beta bursting, and suggest synaptic transmission as a target for future neuromodulatory and therapeutic interventions.

## Materials and methods

### Data collection

The data were previously published, having been acquired by Ben Hunt for the UKMEG partnership [37]. The study was approved by the University of Nottingham Medical School Research Ethics Committee. 75 healthy adult participants were scanned with a 275-channel CTF MEG system. Data were acquired at 1200 Hz inside a magnetically shielded room with 3rd order synthetic gradiometer configuration for reduction of environmental noise. Volunteers were sat upright for the duration of the recording which involved a resting state paradigm whereby participants were asked to stare at a fixation cross for 5 minutes and “think of nothing”. Data were pre-processed by band-pass filtering at 1-150 Hz, and those datasets with excess artifacts (e.g. movement *>* 5 mm) were removed. This resulted in 66 (age 38 *±* 12; 35 female) participants in the final analyses.

### Data analysis

The subsequent analysis pipeline has been previously described in [21]. The cortex was parcellated into 78 AAL regions. An LCMV beamformer was used to compute timeseries at each 4mm voxel in the brain (multiple local sphere head model, covariance matrix computed within the 1-150Hz frequency band for the full 5-minute duration of the recording, Tikhonov regularisation at 5%). For each AAL region, the component voxels were weighted by a Gaussian function of the distance of the voxel from the centre of mass of the regions and summed. This resulted in a single time course of activity for each of the 78 AAL regions in the brain. Each time course was filtered at 1 - 48 Hz and down sampled to 100 Hz. Symmetric orthogonalization was used to reduce special leakage.

A “beta burst state” was identified for each timeseries using a 3-state time-delay embedded hidden Markov model [20] with autocovariance matrix computed with a 230ms time widow. The output of the HMM was a probability value for each state, at each time point (instantaneous probability). The correlation of each state probability timeseries with the beta amplitude envelope was calculated – the state which correlated best with the beta amplitude envelope was identified as the “beta burst state” for each AAL region. Those time points where the probability of being in a “beta burst state” was greater than two thirds were designated as “beta bursts”. Burst parameters (burst duration, burst power, and burst rate) were calculated for each visit to a burst state for every burst across all AAL region in the brain. For a more in-depth description and analysis of the HMM, see Seedat et al. [21] and the Github repository https://github.com/OHBA-analysis/HMM-MAR.

### Mathematical model

We consider the two population next-generation neural mass model of Byrne and Coombes [17, 19], with second order synaptic coupling and gap junction coupling. We introduce an additive stochastic forcing term, *S_a_*, that is the solution to an Ornstein-Uhlenbeck process [38] which we use here to generate low-pass filtered white noise [39]. This low-pass filtering of the noise was done due to beta bursts often occurring alongside alpha-band activity [2]. ( The model may be written as

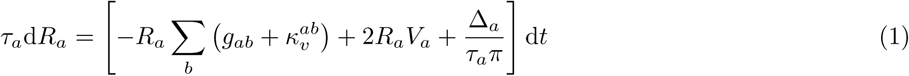

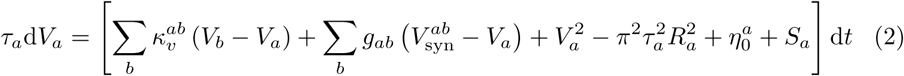

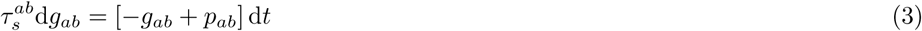

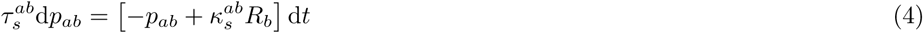

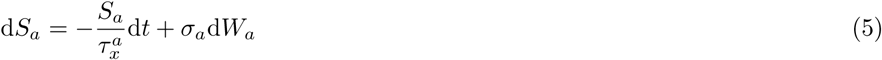

where *a, b ∈ {E, I}*, *R_a_*is the firing rate and *V_a_*is the mean voltage of population *a*, *g_ab_*is the synaptic conductance from population *b* to *a*, *p_ab_*is an auxiliary variable to account for second order synaptic filtering and d*W_a_* is a Wiener process.

We employ a Euler-Maruyama scheme with a step size of Δ*t* = 0.1 ms. We pose the system close to a Hopf bifurcation with oscillations in the beta-band beyond the bifurcation. The stochastic term pushes the system between the stable fixed point and oscillatory regime, creating *bursts* of beta-band activity when the system is beyond the bifurcation point. We restrict to small *σ_a_* to keep the amplitude of the Gaussian noise small.

We fit the model to MEG data which detects signals based on ionic currents (inferred via magnetic field changes). As such, we pass the sum of post-synaptic excitatory and inhibitory currents as the output time-series that we pass to the optimisation algorithm:

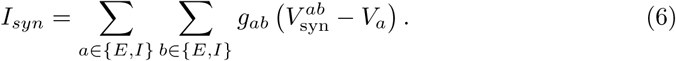

We perform the optimisation using the parameters 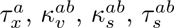.

### Genetic Algorithm

We employ a Genetic Algorithm to find a parameter set for the stochastic neural mass model that has a good fit with the MEG data. Genetic Algorithms are a class of iterative optimisation algorithms first developed by John Holland and colleagues in the 1960s and 1970s that take inspiration from Darwin’s theory of evolution (see [40] for a comprehensive review).

The data we wish to fit the model to are the distribution of burst duration, number of bursts, and burst power from several human MEG resting state data. The total fitness is calculated by taking the average relative absolute difference between the MEG statistics and the model statistics. The fitness function is calculated as

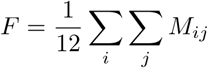

where *M* is the matrix of absolute relative differences of all the target statistics (mean, standard deviation, skew and kurtosis), for each distribution.

### Parameter manipulation simulations

We simulate the model in Python using the built-in Stochastic ODE solver itoint from the sdeint library. For each parameter set, we perform simulations of length 2.5 minutes and repeat each simulation 25 times. We export the synaptic current data to MATLAB and employ the HMM to compute the burst statistics for each trial. The colour plots (Fig. 4 and Fig 5) show the mean burst statistic across the 50 trials.

Initial sweeps were performed across wider parameter windows to identify regions of interest. Once identified, finer-grained parameter sweeps were performed to study these regions in more detail. This process was repeated for locating areas with “restored to healthy” beta burst statistics also.

## Acknowledgments

We acknowledge funding from the UK Quantum Technology Hub in Sensors Imaging and Timing (QuSIT), funded by EPSRC (EP/Z533166/1). We also acknowledge a Medical Research Council (MRC) New Investigator Research Grant (MR/M006301/1) and a MRC Partnership Grant (MR/K005464/1). Funding from EPSRC and MRC (grant number EP/L016052/1) also provided a PhD studentship for ZS through the Oxford Nottingham Biomedical Imaging Centre for Doctoral Training. AB is supported by a Research Ireland Frontiers for the Future Project Grant (22/FFP-P/11547).

## Supporting information

Additional parameter manipulations discussed but not presented in the main next are provided as supplemental information.

**Fig S1.**
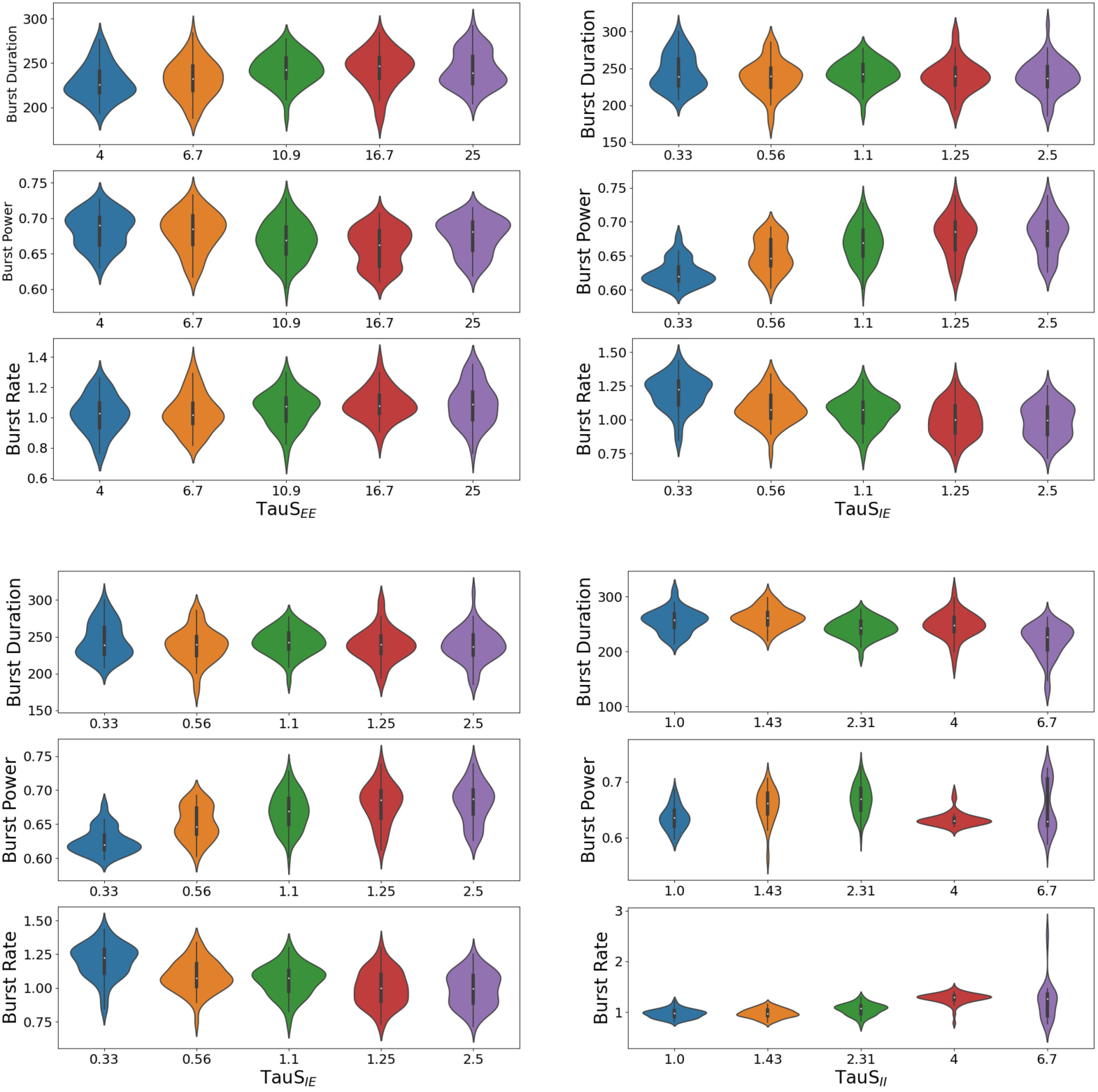
Dependence of the burst statistics on the gap junction coupling strengths. Distribution of the burst duration, burst power and burst rate for various coupling strengths for gap junctions between excitatory neurons *κ^EE^* (top left), between excitatory and inhibitory neurons *κ^IE^* (top right) and *κ^EI^*(bottom left), and between inhibitory neurons *κ^II^* (bottom right).

**Fig S2.**
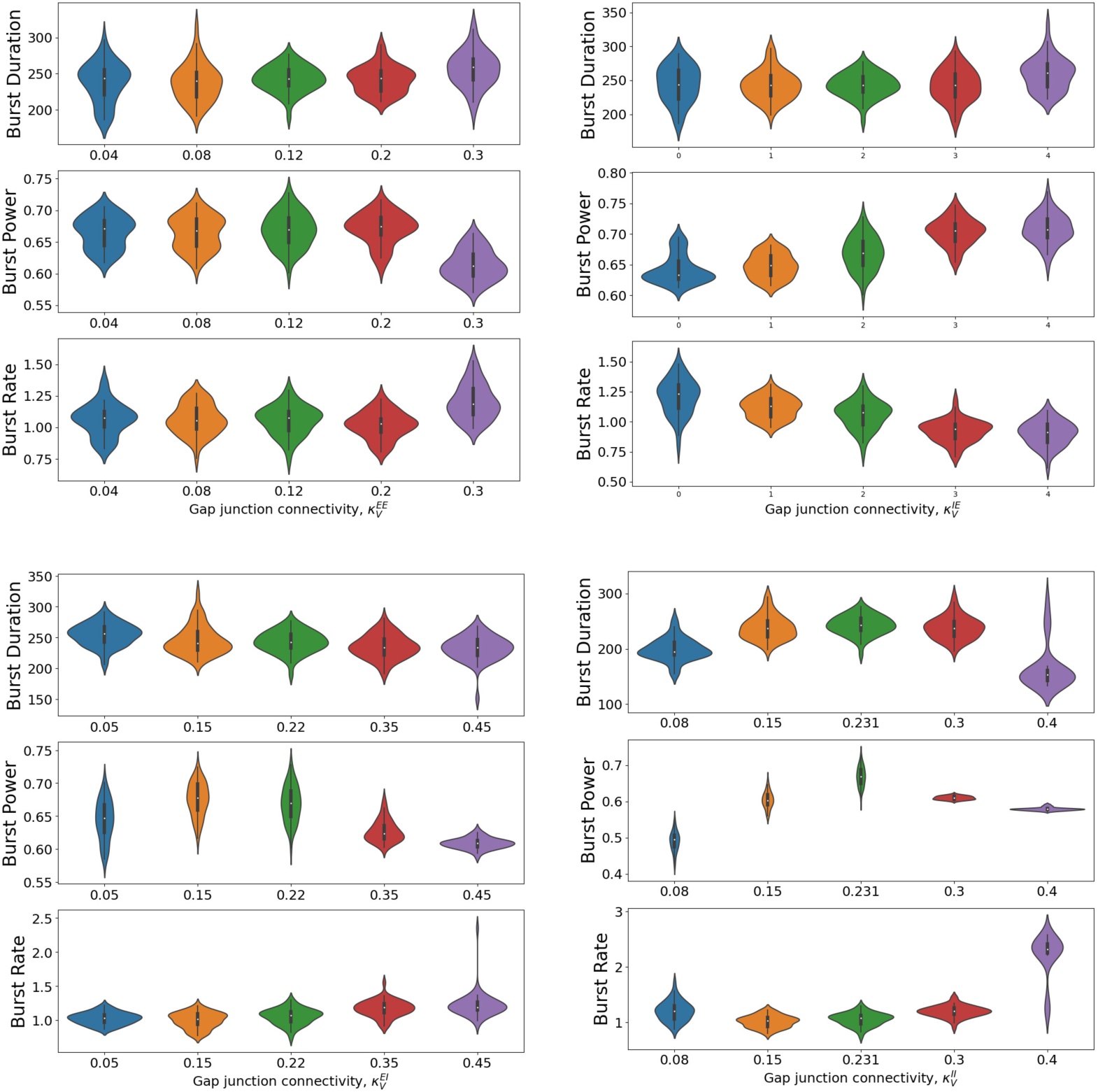
Dependence of the burst statistics on the synaptic time constants. Distribution of the burst duration, burst power and burst rate for various values of the time constant of the synapses from the excitatory population to itself *τ ^EE^* (top left), and to the inhibitory population *τ ^IE^* (top right), as well as the inhibitory population to itself *τ ^II^* (bottom right) and to the excitatory population *τ ^EI^* (bottom left).

**Fig S3.**
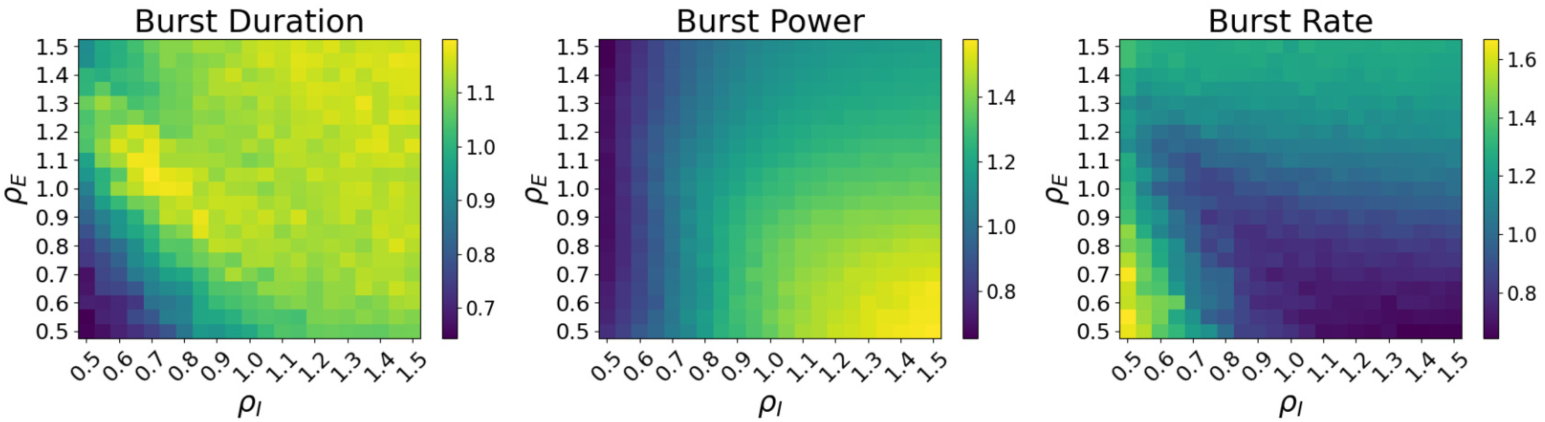
Alternative restoration of healthy activity. Colour plots of the mean burst duration (left), power (middle), and rate (right), relative to their mean values in the healthy case, as the strength of the synaptic connections are varied. The mean background drives are set to the pathological values *η_E_* = 0.8 and *η_I_* = 0.8. All other parameters are set to the optimised values.

